# Molecular reprogramming of adventitial pericytes by a selective MEK inhibitor halts the progression of thoracic aortic aneurysm

**DOI:** 10.1101/2025.02.16.638506

**Authors:** Khaled AK Mohammed, Elisa Avolio, Valeria V Alvino, Eltayeb Mohamed Ahmed, Cha Rajakaruna, Ahmed Ghoneim, Ahmed Elminshawy, Gianni D Angelini, Paolo Madeddu

**Author notes:** **Corresponding authors: Paolo Madeddu,** Emeritus Professor and Honorary Professor of Experimental Cardiovascular Medicine, Bristol Medical School, Translational Health Sciences University of Bristol, Bristol Royal Infirmary, Upper Maudlin Street, BS28HW, Bristol, United Kingdom, **Elisa Avolio**, British Heart Foundation Research Fellow, Bristol Heart Institute, Translational Health Sciences, Bristol Medical School, University of Bristol, Bristol Royal Infirmary, Upper Maudlin Street, BS28HW, Bristol, United Kingdom.

## Abstract

**Aim:** Reconstructive surgery is a life-saving treatment for individuals with advanced thoracic aortic aneurysms (TAAs) at risk of rupture. No effective pharmacological treatments are available to halt aortic dilatation before it reaches this critical point. Unravelling the cellular and molecular pathways involved in TAA formation and expansion is fundamental for identifying potential treatment targets. This study challenged the hypothesis that pericyte dysfunction destabilizes the adventitial vascular niche, hence compromising the ascending aorta’s elastic characteristics. We also investigated whether blocking the mitogen-activated extracellular signal-regulated kinase (MEK) pathway could slow TAA progression in a mouse model.

**Methodology:** Comparative histology and morphometry studies were performed on human ascending TAA and non-aneurismatic control tissues to quantify adventitia vasa vasorum (VV) size and abundance, and pericyte coverage and density. Proliferation, migration, and angiogenesis experiments were used to evaluate the functional phenotype of pericytes before and after blocking the MEK signalling pathway with PD0325901. In a mouse model with moderate TAA, we investigated the therapeutic efficacy of PD0325901 (10 mg/kg/d orally for 14 days) on TAA progression.

**Results:** The histological study of TAA samples demonstrated VV remodelling and reduced pericyte VV coverage due to increased detachment. Cultured TAA pericytes exhibited aberrant behaviour, including increased proliferation, migration, matrix metalloprotease activity, disrupted angiogenic capacity, and altered secretome, and MEK overactivation. PD0325901 restored pericyte contractile phenotype and angiogenic capacity, influencing their secretome, migratory capacity, and matrix formation/degradation equilibrium. In vivo, PD0325901 remarkably decreased aortic dilatation, increased compliance, retained medial elastin content, and reduced adventitial inflammation. No harmful consequences were noted.

**Conclusion:** This work identifies pericyte dysfunction related to MEK overactivation as a major contributor to TAA progression. This suggests that inhibiting the MEK signalling pathway could be a potential treatment option for TAA before surgical intervention becomes necessary.

**Clinical Perspective:** *What is new?:*

- Thoracic aortic aneurysm (TAA) remains a major clinical challenge due to its asymptomatic progression and risk of life-threatening rupture. Our study highlights the crucial role of adventitial pericytes in maintaining vascular homeostasis and preventing aneurysm-related vascular remodeling.
- We demonstrate that dysregulated MEK/ERK signaling drives aortic adventitial pericyte dysfunction, leading to microvascular instability, extracellular matrix degradation, chronic inflammation, and progressive aneurysm expansion.
- Importantly, we show that the clinically available MEK inhibitor, PD0325901, effectively restores pericyte function, preserving adventitial vascular integrity, reducing inflammatory cytokine production, and stabilizing the aortic wall.
- In a preclinical mouse model, PD0325901 significantly attenuated aneurysm growth, improved aortic wall compliance, and prevented maladaptive vascular remodeling.

*What Are the Clinical Implications?:*

- Our findings provide strong translational evidence supporting MEK inhibition as a promising therapeutic strategy to halt TAA progression and enhance aortic wall resilience, offering a potential medical alternative to delay or prevent surgical intervention.

## Introduction

Thoracic aortic aneurysms (TAA), particularly those located in the proximal aorta, are life-threatening vascular disorders characterized by progressive pathological dilatation and weakening of the aortic wall, ultimately leading to dissection or fatal rupture ^1^. Despite advancements in imaging and surgical techniques, the lack of pharmacological interventions to prevent aneurysm progression highlights a significant unmet clinical need ^2^. The current management of TAA is mainly achieved through surgical correction ^3^. This approach is recommended when the aorta reaches a size (5-5.5 cm) where the risk of complications (i.e., acute dissection or fatal rupture) equals or exceeds the operative risk ^4^. However, this criterion depends on the underlying pathology, growth rate, family history, and individual patient ^1^. Many patients remain in the grey zone, where the aortic diameter stays below the hinge point for surgical intervention. Pharmacological treatments recommended for this grey stage limit the impact of risk factors but are not targeted to pathogenic mechanisms ^5^.

While the pathogenesis of TAA has been traditionally associated with medial layer degeneration, emerging evidence has shifted the focus to adventitia as a critical player in the initiation and progression of aneurysm pathology ^6^. Within the adventitia, a vascular network, the vasa vasorum (VV), provides vital nutrition to large arteries and veins. The VV also represents a dynamic unit capable of developing homeostatic remodeling responses to chemical and mechanical changes and injuries at the luminal side of the blood vessel ^6,7^. The anatomy of the parent vessel dictates the need for VV, and, in turn, the VV abundance and organization oversee vessel health and homeostasis. VV are prominent in the aortic arch, in line with the greater metabolic needs of this large artery ^8^, and typically penetrate from the adventitia into the media, at variance with VV of other arteries ^9^. Experimental studies collected using animal surgical models revealed the impact of VV loss on aortic function ^10^. However, evidence of whether VV remodeling participates in the progression of AA remains limited.

Several groups reported that pericyte-like cells expressing progenitor markers and capable of clonogenic expansion reside near the adventitial VV of human adult and fetal aorta ^11,12^ and vena saphena ^13^. Seminal studies suggest that adventitial pericytes (APCs) maintain aortic wall stability but also contribute, when dysfunctional, to adverse vascular remodeling ^13,14^. At the organismal level, pericyte loss is associated with vascular fragility, altered barrier function, and fibrotic remodeling ^15,16^, all of which are hallmarks of TAA. Despite their recognized importance in vascular biology, the role of APCs in TAA pathogenesis remains poorly understood.

The mitogen-activated protein kinase (MAPK)/extracellular signal-regulated kinase (ERK) signaling pathway is known to regulate critical cellular processes such as proliferation, migration, and differentiation ^17^. Dysregulation of this pathway has been implicated in various vascular diseases, including aneurysms. Overactivation of ERK signaling has been observed in TAA, suggesting its involvement in the pathological remodeling of the aortic wall ^18–21^. However, the precise molecular and cellular mechanisms linking MEK/ERK dysregulation to adventitial remodeling and APC dysfunction in TAA remain unclear. Targeting this pathway using selective inhibitors might offer a promising therapeutic approach to stabilize the aortic wall and mitigate aneurysm progression.

This study investigates the role of APCs in VV remodeling and TAA progression. Using a combination of human tissue histological analyses, in vitro functional and molecular assays, and an established murine model of aortic aneurysm, we provide the first evidence that dysfunctional APCs play a central role in adventitial remodeling, likely due to upregulation of the MEK/ERK pathway. Furthermore, we demonstrate the therapeutic potential and safety of PD0325901, a selective MEK inhibitor (MEKi), in restoring APC function and preventing aneurysm progression.

## Material and Methods

### Ethics and sample collection

This study complies with the guidelines of the Declaration of Helsinki. Human aortic samples were collected from Bristol Heart Institute, Bristol, UK, in compliance with the waste tissue ethical protocol (REC reference: 06/Q2001/197). **Supplementary Tables 1 and 2** provide data on patients and aortas collected for this study. Each collected sample (if sufficient) was divided into two parts, one used for histological studies and the other for APCs isolation and expansion.

### Histology and Immunohistochemistry (IHC)

Freshly collected samples were immediately fixed by incubation in 4% w/v paraformaldehyde (Sigma-Aldrich) at 4°C for 24 hours. Following fixation, the samples were processed for histology, embedded in paraffin, and sectioned into 5-μm sections, which were mounted onto microscopy slides (Thermo Fisher). Slices were then dewaxed, rehydrated, and subjected to heat-retrieval using citrate buffer (0.01 M, PH 6, for 30 min at 98°C). For morphometric analysis, sections were stained with Hematoxylin and Eosin (H&E) and Van Gieson stain. For IHC, sections were incubated with 5% v/v donkey serum to prevent non-specific antibody binding and minimize background signal. Sections were then incubated overnight at 4°C with primary antibodies **(Supplementary Table 3)**. Samples were incubated with donkey Alexa-Flour 488, 568 or 647-conjugated secondary antibodies (all from ThermoFisher) at room temperature for 1 hour. This was followed by nuclear staining with DAPI and mounting with Fluoromont-G (Thermo Fisher Scientific, # 00-4958-02). Imaging was done using an Axio Observer Z1 fluorescent microscope (Zeiss).

### APC isolation, culture and expansion

The explant outgrowth method was employed to isolate APCs ^22^. Aortic adventitial samples were cut into small fragments (approximately 3 mm^2^) and seeded onto tissue culture flasks or 6-well plates. These fragments were cultured in full endothelial growth media (EGM2) supplemented with 5% fetal bovine serum (FBS) and maintained in an incubator set at 37°C with 5% CO_2_. Media was replenished every 48 hours, and cellular outgrowth was observed sprouting from the adventitial explants after 5-7 days of incubation, showing the characteristic shape of pericytes **(Supplementary Figure 1A&B)**. Once the cells reached confluence, the tissue explants were removed, and cells were detached using Acutase solution (Sigma, A6964) for subculturing and expansion. All experiments were conducted using cells between passages 3 and 6.

### Viability, proliferation, and doubling time

A 6-well tissue culture plate was utilized to seed 30,000 cells per well which were then maintained in full EGM2 media within an incubator set at 37°C and 5% CO2. Cells were detached and counted on days 4, 5, 6 and 7 to generate a growth curve. Cell counting was performed using trypan blue (Lonza, # 17-942E) and a Bürker counting chamber (Hawkslev). The number of viable cells was recorded to calculate the cell doubling time. Cell viability was assessed with a Calcein-AM/EthDIII staining (Biotium, Cat# 30002) according to the manufacturer’s instructions. Proliferation was assessed using the Click-iT EdU Cell Proliferation Kit (C10337 - ThermoFisher Scientific) according to the manufacturer’s instructions and an anti-Ki67 antibody **(Supplementary table 4).**

### Immunocytochemistry (ICC)

A 96-well black plate was used to seed 5,000 cells per well followed by incubation in full EGM2 media at 37°C in a 5% CO2 incubator. After 48 hours, cells were fixed with 4% v/v PFA, permeabilized and incubated with 5% FBS for 1 hour to minimize non-specific antibody binding. Subsequently, cells were incubated overnight at 4°C with primary antibodies **(Supplementary Table 4)**. After washing with PBS, cells were incubated with donkey Alexa-Flour 488, 568 or 647-conjugated (all from ThermoFisher) secondary antibodies at room temperature in the dark for 1 hour. Nuclear counterstaining was performed using DAPI, and cells were preserved using Fluoromont-G (Thermo Fisher Scientific, # 00-4958-02). Imaging was conducted using an inverted Axio Observer Z1 Fluorescence Microscope (Zeiss).

### Analysis of the secretome by ELISA

After cultured cells reached 80-90% confluence, media was removed, and cells were rinsed with DPBS. Cells were then incubated with basal EBM2 media (i.e., serum-free and growth factors-free media) for 48 hours. The conditioned media was collected, centrifuged to remove floating cells and debris, and stored at -80°C until analysis. Cells were detached and counted to normalize the levels of secreted factors versus the number of cells. ELISA kits (R&D Systems) were used to quantify secreted angiogenic factors, including Angiopoietin-1 (ANGPT-1) (#DY923) and Angiopoietin-2 (ANGPT-2) (#DY623) and thrombospondin-1 (TSP-1) (#DY3074), extracellular matrix proteins Fibronectin (#DY1918) and Collagen 1-alpha (#DY6220), and pro-inflammatory factors Interleukin-6 (IL-6) (#DY206) and Monocyte Chemoattractant Protein-1 (MCP-1) (#DY279). The levels of secreted factors were normalized against the final total cell number and the time. Absorbance was measured using a GloMax Discover System.

### Angiogenic tube formation assay

The ability of isolated APCs to interact with and support the angiogenic tube networks formed by aortic endothelial cells (AoECs) was assessed. APCs were co-cultured with AoECs at a 2:5 ratio in EBM2 media supplemented with 2% FBS. Cells were plated on a thin layer of ECM gel (Sigma, # 6909) within the wells of angiogenesis slides (Ibidi, # 81506). Cells were incubated for 6 hours at 37°C in a 5% CO2 incubator before being imaged using a brightfield microscope (Leica). Experiments were performed in triplicate, and the total tube length per image was measured using ImageJ Software.

### RT-qPCR

Upon confluence, cells were lysed using Qiazol (Qiagen, # 1023537) and total RNA was extracted using RNeasy Plus kit (Qiagen, # 74134). After quantification, equal levels of RNA were reverse transcribed into cDNA using RNA to cDNA kit (Applied Biosystems, # 4387406). Realtime quantitative PCR (qPCR) was then used to quantify the expression levels of different mRNAs using TaqMan Master Mix using a QuantStudio 5 System (Applied Biosystems). The relative gene expression was calculated using the 2^-ΔΔCt^ mathematical method ^23^. The list of TaqMan probes used for qPCR analysis is provided in **(Supplementary Table 5)**.

### Western blotting

Confluent cells were rinsed with DPBS and lysed with RIPA buffer (Sigma, # R0278) supplemented with phosphatase and protease inhibitors (Sigma Aldrich). Protein concentration was quantified using the BCA Protein Assay kit (Thermo Scientific, # 23250). Equal amounts of total protein (10-20 μg) from each sample were loaded onto TGX 7.5% gels (Bio-Rad, # 4561024), and separated via electrophoresis using running buffer (Invitrogen, # LC2675) for 90 minutes at 110-150 V (PowerPac HC, Bio-Rad). Proteins were transferred to PVDF membranes (Bio-Rad, #1620177) using transfer buffer (Novex, # LC3675) at a constant current of 0.25 A for 60 minutes. Membranes were washed and blocked for 2 hours in a solution of 5% globulin-free bovine serum albumin (Sigma, # A3059) in TBST solution (i.e., TBS (Bio-Rad, # 1706435) + 0.05% v/v Tween-20 (Sigma, # P1379)). Membranes were then incubated overnight at 4°C with primary antibodies, including ERK 1/2 (Cell Signaling, # 4695) and P-ERK 1/2 (P44/42 MAPK, Cell Signaling, # 4370) (**Supplementary Table 6)**. The following day, membranes were washed and incubated at room temperatures for 1 hour with horseradish peroxidase (HRP)-conjugated anti-rabbit (GE Healthcare, # NA934V) secondary antibody. After washing, protein bands were visualized using a Chemi-Doc MP system (Bio-Rad) following enhancement of chemiluminescence signals with ECL detection Reagents (Cytiva, # PRN2232) for 5 minutes. Band densitometry was analyzed using ImageJ software. The densitometry values for the target proteins were normalized versus β-Tubulin loading controls.

### Migration Assay

A wound closure migration assay was conducted to evaluate the migratory capacity of isolated APCs. In this assay, 15,000 cells were seeded into each well of a 96-well plate. Once the cells reached confluence, a scratch was created in the center of the well using a 20 μL pipette tip. Cells were then incubated in basal (i.e., serum- and growth factor-free) EBM2 media supplemented with hydroxyurea (2 mM) to inhibit proliferation. Human recombinant platelet-derived growth factor beta (PDGF-BB, Peprotech, 20 ng/mL) was used to stimulate cell migration. Absence of growth factors was used as control. Images of the scratch area were taken at 0, 6, 8 and 10 hours using an inverted microscope (Leica). The scratch/wound area was quantified using ImageJ software.

### Gelatinase assay

The gelatinase activity of the isolated APCs, specifically MMP-2 and MMP-9, was evaluated using a commercial kit (Sigma-Aldrich, # MAK348) according to the manufacturer’s protocol. In this assay, 100,000 cells were seeded per well in a 6-well culture plate. Once confluent, cells were lysed using the provided cell lysis buffer. The lysate was then centrifuged at 16,000x*g* for 10 minutes at 4°C, and the supernatant was collected into an Eppendorf tube on ice. Protein concentration in the supernatant was determined using the BCA Assay kit (Thermo Scientific, # 23250). Equal protein concentrations (20 μg) from each sample were loaded into the wells of 96-well black plate. Reconstituted gelatin substrate was added to each well, and fluorescence was measured using a fluorescence microplate reader (GloMax). Gelatinase activity was expressed in units per milligram (U/mg) where one unit (U) is defined as the amount of gelatinase required to cleave the gelatine substrate per minute.

### PD0325901 treatment on cultured APCs

To investigate the *in vitro* effects of MEK/ERK signaling inhibition, AA-APCs were treated with the selective MEK inhibitor PD0325901 (Sigma-Aldrich, #PZ0162). For this study, cells were divided into two groups (i.e., vehicle and MEKi). Upon reaching confluence, the MEKi group was treated with 250 nM PD0325901 for 10 days, with media being replaced every 48 hours. The vehicle group was treated with an equivalent dilution of the solvent (DMSO) under identical conditions, including the same frequency of media replacement. At the end of the 10-day treatment period, cells were used for antigenic and functional characterization using the methods described above.

### In vivo study on a mouse model of aortic aneurysm

The study was covered by a license from the British Home Office (PP1377882) and complied with the EU Directive 2010/63/EU. Procedures were carried out according to the principles in the NIH Guide for the Care and Use of Laboratory Animals (National Academies Press, 2011). Termination was conducted according to humane methods outlined in the Guidance on the Operation of the Animals (Scientific Procedures) Act 1986 Home Office (2014). The report of results is in line with the ARRIVE guidelines.

The *in vivo* effects of PD0325901 were evaluated in a mouse model of Angiotensin-II (ANG-II) induced aortic aneurysm as previously described ^24,25^. In brief, basal echocardiography was performed before osmotic minipumps (Model 2004; ALZET) were implanted (under recovery anesthesia using inhaled isoflurane 2-3%) in fourteen 8-week-old male C57BL/6J mice (Charles River) for continuous subcutaneous ANG-II infusion (rate = 1000 ng/kg/min, Enzo Life Sciences, UK). Seventy-two hours post-implantation, mice were randomly assigned to two groups: the treatment group received oral PD0325901 at a dose of 10mg/kg per day (n=7), and the vehicle group received oral DMSO (n=7) for 14 days. The drug and control solvent were embedded in strawberry-flavored sugar-free jelly as previously described^26^. Final echocardiography was conducted before humane euthanasia, followed by blood and organ collection. A control group (n=5) that did not receive ANG-II or DMSO/PD0325901 treatment was included for reference control. **Supplementary Table 7** shows echocardiography parameters for each group.

Harvested aorta samples were fixed in 4% PFA at 4°C overnight, immersed in 30% w/v sucrose in PBS overnight and embedded in OCT for cryopreservation and histological analyses. 5-μm thick frozen aorta sections were post-fixed and permeabilized with ice-cold acetone (VWR) for 5 minutes at -20°C and let air dry for 30 minutes. Sections were rehydrated with PBS for 10 minutes before proceeding with histological and immuno-staining as described above.

For western blotting, freshly frozen aorta samples were homogenized in RIPA buffer supplemented with proteases and phosphatases inhibitors using gentle MACS M tubes (Miltenyi). Tissue lysates were centrifuged at 10,000 *g*, at 4°C, for 15 minutes and processed as described above for detection of P-ERK / ERK protein levels.

### Statistical analysis

Statistical analysis was conducted using Prism GraphPad software. The D’Agostino-Pearson and Kolmogorov-Smirnov normality tests were used to check for data normal distribution. Continuous normally distributed variables were compared using the Student’s t-test (2 groups) or 1-way ANOVA (multiple groups). Nonparametric tests, including the Mann-Whitney (2 groups) and Kruskal-Wallis (multiple groups), were used to compare data not normally distributed. Post hoc analyses included Tukey’s and Dunn’s comparison tests. For in vivo studies, 2-way ANOVA was used to compare the mean differences between groups, followed by Sidak’s test. Baseline and final echocardiography parameters in the same animal were compared using paired tests; for all other analyses, unpaired tests were applied. Data are presented as means+/- SEM (i.e., Standard error of the mean). Statistical significance was defined as a *p-value* less than 0.05.

## Results

### Vasa vasorum remodeling and APC detachment in human TAA samples

Analysis of H&E and Van Gieson staining revealed notable VV remodeling in AA samples compared to controls, appearing larger in size with wider lumens and thicker walls **(Figure 1A&B)**. The morphometric evaluation demonstrated a significant reduction in VV density within the adventitia of TAA samples compared to controls (1.7±0.3 vs. 5.5±0.6 per microscopic field) **(Figure 1C)**. Moreover, quantitative analysis revealed an approximately 8-fold increase in the lumen area of TAA-VV compared to control-VV (1039±163 vs. 126±25 μm^2^) **(Figure 1D**), possibly representing a compensatory response to VV rarefaction. Normalizing VV medial area to the lumen diameter showed that TAA-VV have significantly thicker walls (19.8 ± 1.5 μm^2^/μm) compared to controls (10.4 ± 0.9 μm^2^/μm) **(Figure 1E)**, which can be considered an eccentric hypertrophy.

**Figure 1.**
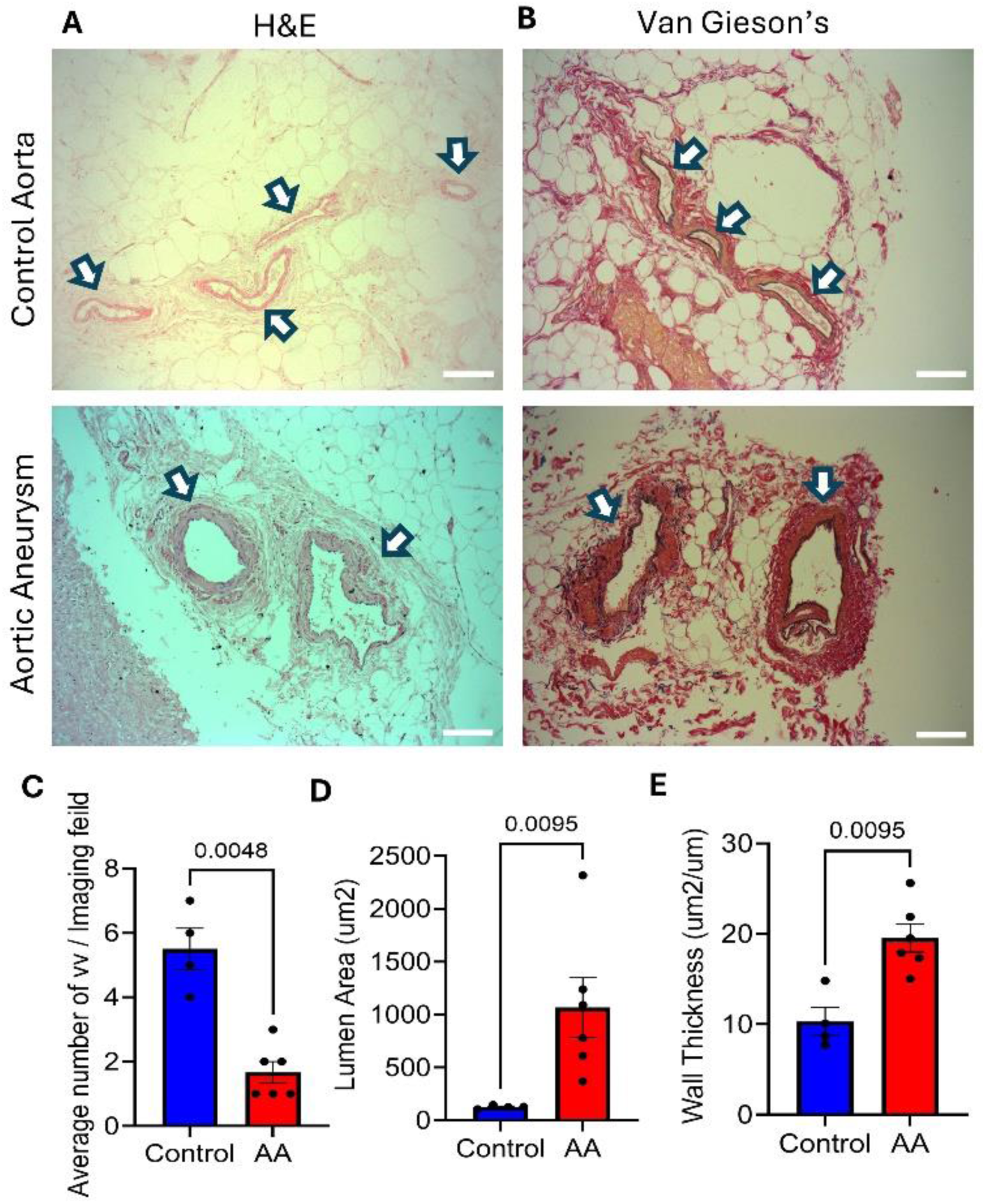
Histological analysis of human adventitia samples. **(A)** Haematoxylin & Eosin (H&E) and **(B)** Van Gieson’s stainings. Scale bar = 50 µm. White arrows point to VV vessels. Images are representative of n=1 sample of each group. **(C-E)** Results of statistical analysis of morphometric measurements for **(C)** VV density, **(D)** VV lumen area, and **(E)** VV wall thickness. N = 4 control and 6 TAA. Data are presented as individual values and means ± SEM. Analysis by unpaired Mann-Whitney test.

APCs surrounding adventitial VV were identified as platelet-derived growth factor receptor beta (PDGFRβ)^+^/CD34^+^/CD31^-^ cells **(Figure 2A)**. IHC analysis revealed a significant reduction in APC coverage of VV in TAA samples compared to the controls **(Figure 2B)**. In control samples, APCs covered 72.7±2.3% of VV surface area whereas this coverage was significantly reduced to 49.4±1.7% in TAA samples **(Figure 2C)**. To further investigate VV remodeling, correlation analysis indicated a negative relationship between VV perimeter and APC coverage, where larger perimeters corresponded to reduced APC coverage **(Figure 2D)**. Interestingly, while APCs in control samples tend to preserve their intimate contact with VV, we observed APC detachment from their peri-VV location in TAA samples, with APCs appearing denser in the proper aorta adventitia (**Figure 2E&F**). This suggests that AA-VV rarefaction and enlargement are associated with progressive APC detachment and migration away from the original perivascular site.

**Figure 2.**
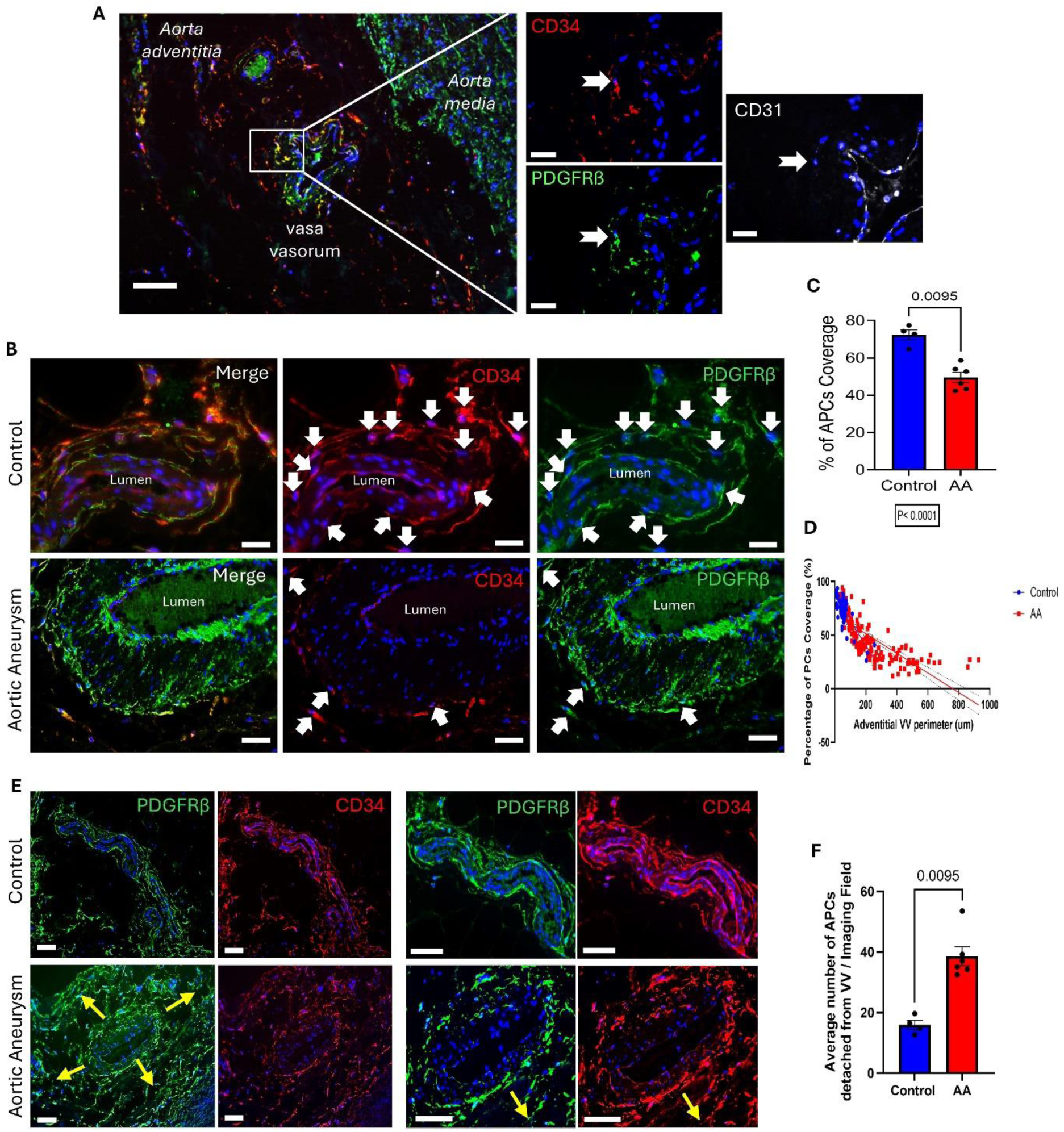
*In situ* characterization of APCs in human aorta adventitia using IHC. **(A)** Peri-VV-associated APCs are characterized by expression of PDGFRβ and CD34 and lack of CD31. **(B)** Immunofluorescence images showing the different coverage of VV with PDGFRβ+/CD34+ APCs (white arrows) in the two groups. PDGFRβ = Platelet-Derived Growth Factor Receptor beta (green), CD= Cluster of Differentiation (red). DAPI (blue) labels the nuclei. Scale bar = 50µm. **(C)** Bar graphs showing a significant decline in APC coverage in TAA samples compared to controls. N= 4 control and 6 TAA. Data are presented as individual values and means ± SEM. Analysis by unpaired Mann-Whitney U test. **(D)** Correlation study between the APC coverage and VV perimeter. The study showed a significant difference between the slops (p<0.0001) by simple linear regression test. **(E)** Immunofluorescence images showing detachment of APCs from VV into the adventitia of TAA samples, yellow arrows show the direction of detachment. Scale bar = 50 µm. **(F)** Bar graphs reporting the quantification of APCs detaching from VV. PDGFRβ = Platelet-Derived Growth Factor Receptor beta (green), CD= Cluster of Differentiation (red). DAPI (blue) labels the nuclei. N= 4 control and 6 TAA. Data are presented as individual values and means ± SEM. Analysis by unpaired Mann-Whitney U test.

### APC isolation and antigenic characterization

Using the explant outgrowth method, we isolated APCs from aneurysm and control aorta adventitia samples to better characterize functional properties. Cells sprouting from the adventitial explants were observed after 5-7 days of incubation **(Suppl. Figure 1A)**. The isolated cells exhibited the characteristic morphology of pericytes, appearing as spindle-shaped cells with small central nuclei **(Suppl. Figure 1B)**, consistent with previous descriptions ^13,22^. Antigenic characterization of sprouting cells at passage zero confirmed expression of typical APC markers CD34, PDGFRβ, and neural-glial antigen 2 (NG2) and lack of the fibroblast marker PDGFRα and endothelial cell markers CD146 and CD31, as previously reported by our group **(Suppl. Figure 1C-E)** ^13,26^.

Antigenic profiling of expanded cells at passage 4 revealed no differences between APCs isolated from TAA samples and controls. Cells in both groups demonstrated strong positive expression of PDGFRβ and NG2, with negligible expression of CD31 and CD146, smooth muscle alpha-actin (α-SMA), and PDGFRα **(Figure 3A&B)**. As expected, APCs downregulated CD34 during expansion in vitro ^13,26^. Furthermore, cells from both groups showed positive expression of progenitor cell markers NANOG, OCT-4, SOX-2, and GATA-4 **(Figure 3C&D)**, confirming their progenitor status and differentiation potential.

**Figure 3.**
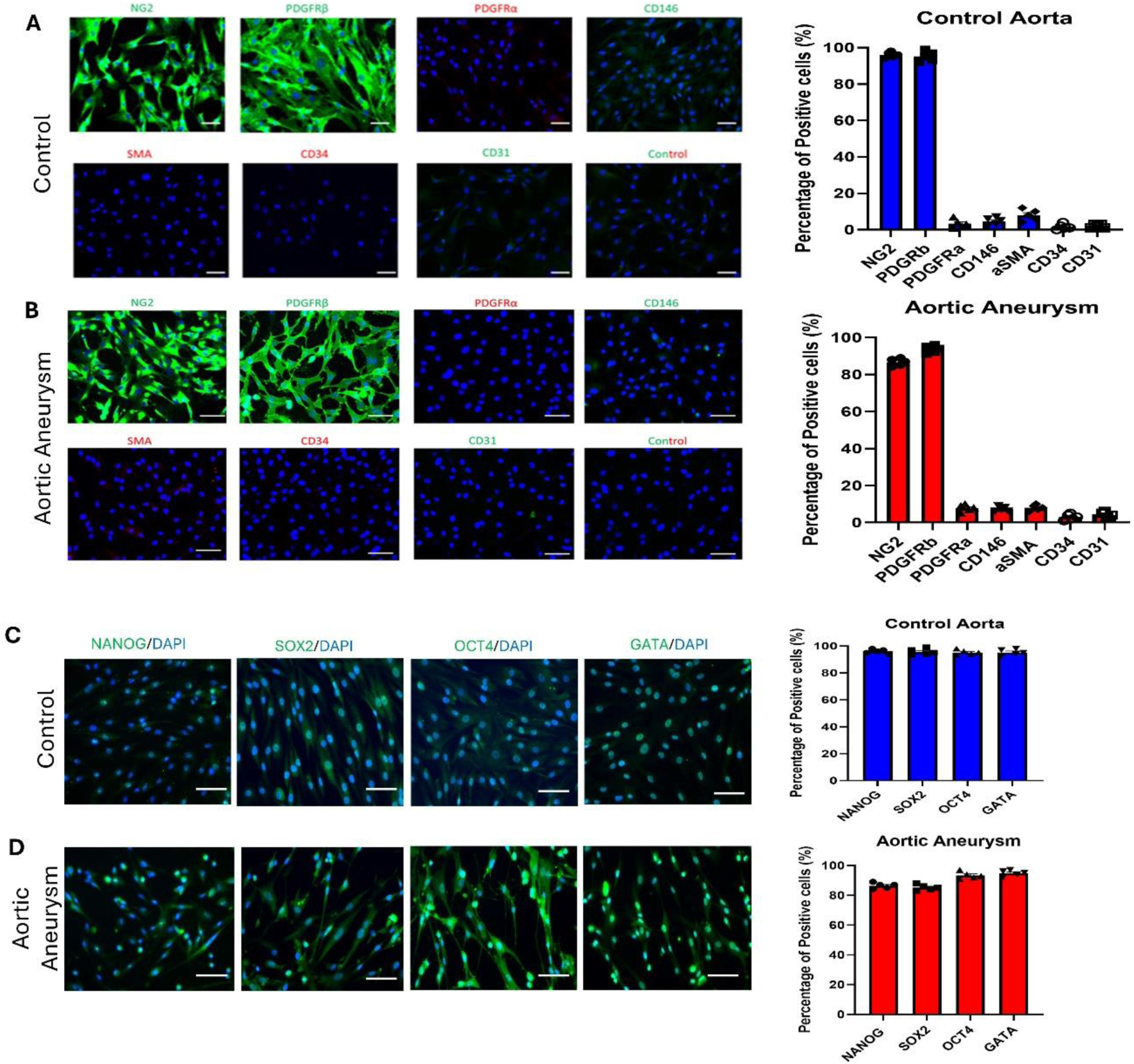
Antigenic characterization of isolated APCs. **(A&B)** Immunofluorescence images and bar graphs showing antigenic profile of **(A)** Control-APCs, and **(B) T**AA-APCs at passage 4 of culture. Both groups show strong positivity to pericyte markers NG2, PDGFRβ, and negativity to tested EC, VSMC and fibroblast markers. NG2=Neural-Glial Antigen 2, PDGFRβ/α= Platelet-Derived Growth Factor Receptor beta / alpha, CD= Cluster of Differentiation, SMA= Smooth Muscle Actin. N = 5. Images are representative of 1 sample from each group. DAPI labels nuclei in blue. Scale bar= 50µm. **(C-D)** Immunofluorescence images and graphs showing progenitor markers expression for **(C)** Control APCs, and **(D)** TAA-APCs. NANOG = homeobox NANOG protein, SOX2 = sex determining Y region-box 2, OCT4 = octamer binding transcription factor 4. N = 5. Images are representative of 1 sample from each group. Scale bar= 50µm. All data are presented as individual values and means ± SEM.

### APCs isolated from aortic aneurysms have impaired molecular and functional properties

TAA-APCs are characterized by altered cellular behavior compared to control cells. Analysis of cell growth revealed TAA-APCs have higher proliferation rates and shorter doubling times (average 72 hours in TAA group vs 106 hours in the control group) **(Figure 4A)**. Additionally, TAA-APCs exhibit enhanced motility, as demonstrated by a wound-healing migration assay in which cells were stimulated with PDGF-BB **(Figure 4B)**. This highly migratory phenotype, in line with APC detachment from VV previously observed in situ, suggests a pathological shift that may exacerbate VV remodeling in TAA.

**Figure 4.**
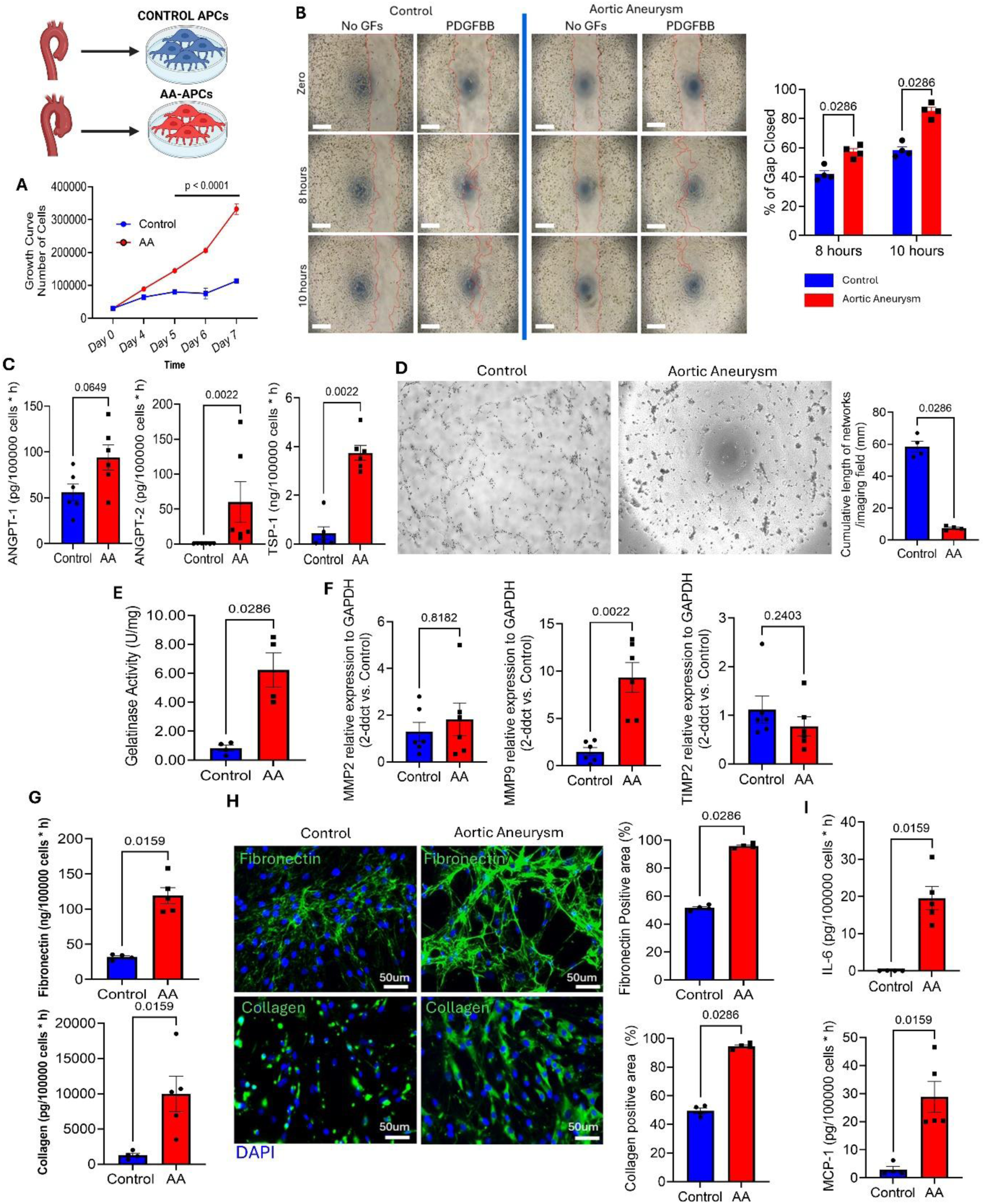
TAA-APCs show impaired functional and molecular properties. **(A)** Growth curve showing hyper-proliferative status of TAA-APCs compared to controls. N= 5. Analysis by 2-way ANOVA. **(B)** Wound closure migration assay comparing non stimulated and PDGFBB-stimulated cells. Images were snapped after 8 and 10 hours and compared with time zero. Bar graphs report the percentage of gap closed. N = 4. PDGFBB = platelet-derived growth factor-BB. **(C)** Secreted angiogenic factors in APC conditioned media analysed by ELISA. N = 6. **(D)** Angiogenic tube formation assay of APCs co-cultured with human aortic ECs (AoECs). Images are representative of 1 sample from each group. N = 4 APCs, n=1 AoECs. **(E)** Gelatinase activity fluorescence assay. Results are presented as U/mg. N = 4. **(F)** Expression of *MMP2*, *MMP9* and *TIMP-1* transcripts as assessed by RT-qPCR. N = 6. **(G)** Secreted ECM proteins fibronectin and collagen in APC conditioned media analysed by ELISA. N = 5 AA-APCs and 4 controls. **(H)** Immunofluorescence images and bar graphs showing fibronectin and collagen expression (Green Fluorescence) in APCs. DAPI labelled nuclei in blue. N = 4 from both groups. **(I)** Secreted pro-inflammatory cytokines, IL-6 and MCP-1 in APC conditioned media analysed by ELISA. N = 5 TAA-APCs and 4 controls. All data are presented as individual values and means ± SEM. In (B-I) analysis by unpaired Mann-Whitney U test.

TAA-APCs also exhibit an impaired angiogenic profile, evidenced by significantly higher secretion of anti-angiogenic factors, including ANGPT-2 and TSP-1, which cause vascular regression **(Figure 4C)**. Moreover, an angiogenic tube formation assay showed impaired APC ability to support the formation of angiogenic networks in co-culture with aortic endothelial cells (AoECs) on Matrigel **(Figure 4D)**. The diminished angiogenic capacity of TAA-APCs reflects a crucial defect that disrupts vascular homeostasis in the aortic adventitia, a pathogenic hallmark of TAA.

TAA-APCs showed elevated gelatinase activity **(Figure 4E)**, associated with increased *MMP9* gene expression **(Figure 4F)**, indicating enhanced extracellular matrix (ECM) degradation. Simultaneously, TAA-APCs secrete higher levels of ECM proteins, including fibronectin and collagen, contributing to abnormal ECM deposition as confirmed by both ELISA and ICC **(Figure 4G&H)**. These changes are accompanied by elevated levels of pro-inflammatory cytokine IL-6 and MCP-1 **(Figure 4I)**, further driving ECM degradation and inflammation. Together, these alterations, indicative of a shift of APCs toward an anti-angiogenic, pro-fibrotic, and pro-inflammatory phenotype, are compatible with immunohistochemical findings and indicative of destabilizing properties resulting in adverse remodeling of the adventitia VV.

### Altered TAA-APC properties are driven by MEK/ERK signaling and restored by its inhibition

We next investigated molecular alterations underlying TAA-APC dysfunction. We previously showed that ERK1/2 signaling controls cardiac pericyte function ^26^. Here, TAA-APCs showed significantly increased levels of phosphorylated ERK1/2, indicating aberrant activation of this pathway **(Figure 5A)**. Therefore, we used a selective MEKi, PD0325901, to modulate the MEK/ERK signaling *in vitro*. PD0325901, at a dose of 250 nM, successfully prevented ERK1/2 phosphorylation/activation in TAA-APCs **(Figure 5B)**, and inhibited – as expected – the translocation of P-ERK to the nuclei **(Figure 5C)**. Conditioning of TAA-APCs with PD0325901 for 10 days did not affect cell viability, as assessed with the Calcein/EthD-III assay **(Figure 5D).** Treated cells showed morphological changes, including increased cell size and the appearance of intracellular striations **(Figure 5E)**. Additionally, while retaining the expression of pericyte/VSMC markers PDGFRβ and NG2, PD0325901-treated cells expressed typical VSMC contractile proteins, including α-SMA, Calponin, smooth muscle myosin heavy chain (SM-MHC), Smoothelin B and Transgelin **(Figure 5F).**

**Figure 5.**
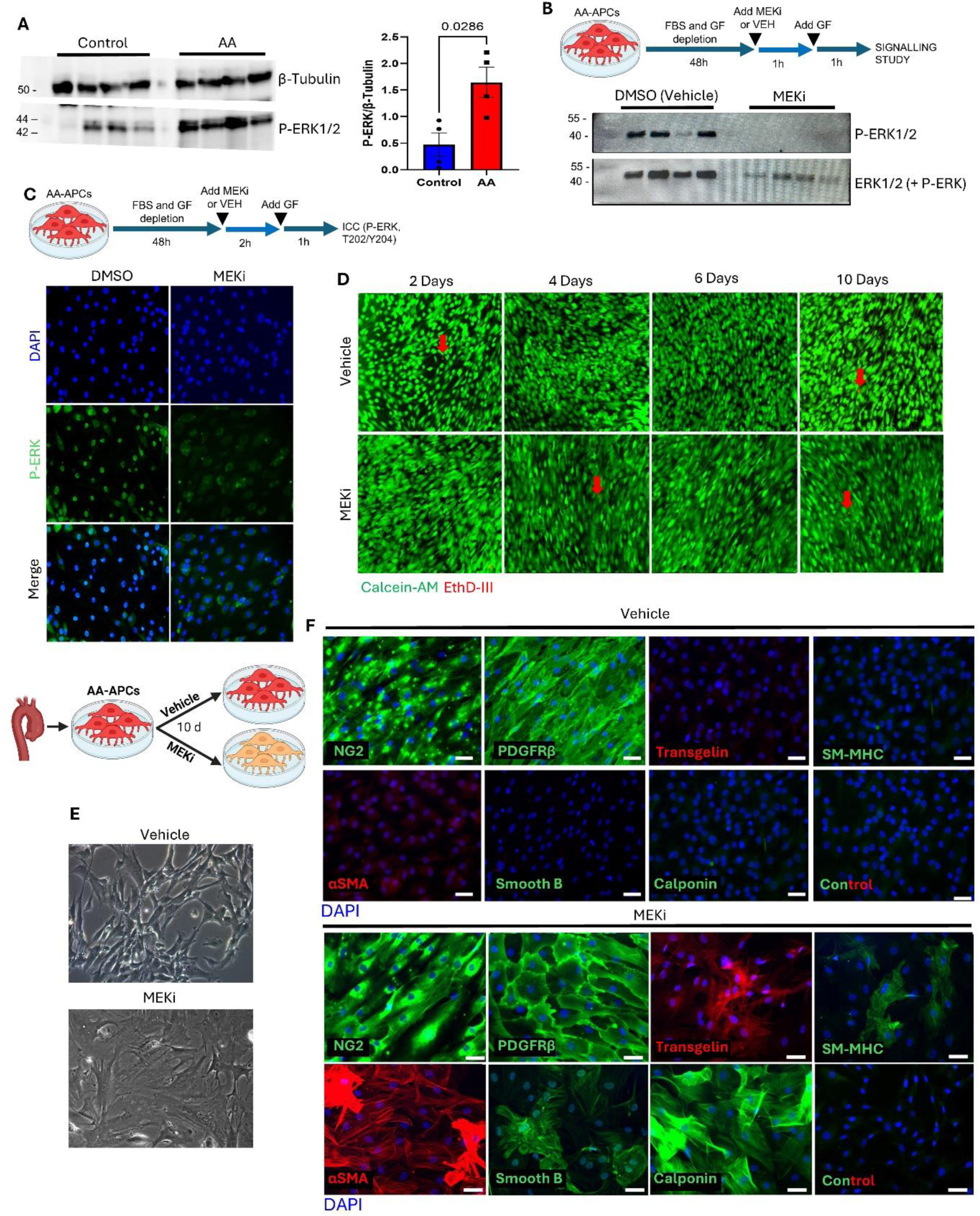
Higher P-ERK levels in TAA-APCs can be regulated by PD0325901. **(A)** Western blot analysis showing TAA-APCs express higher levels of P-ERK compared to controls. N= 4. Data are presented as individual values and means ± SEM. Analysis by unpaired Mann-Whitney U test. **(B)** PD0325901 (MEKi) in a dose of 250nM successfully inhibited ERK phosphorylation/activation in-vitro. N = 4. **(C)** Immunofluorescence images showing MEKi treatment inhibited P-ERK (green) nuclear translocation. DAPI labels nuclei in blue. N = 2. Images are representative of 1 sample. In **(B&C)** APCs were incubated for 48h with a growth factor (GF)-free and serum-free media to switch off molecular signalling, followed by incubation with either the MEKi or its vehicle for 1h, finally followed by stimulation with a GF mix to induce ERK phosphorylation. **(D)** Viability assay in APCs using Calcein-AM (green, viable cells) and EthD-III (red, dead cells). N = 3. Images are representative of 1 sample. **(E)** Brightfield optical images showing morphological changes of TAA-APCs after a 10-day treatment with the MEKi. Cells became larger in size with appearance of intra-cellular striations. **(F)** Immunofluorescence images showing the antigenic profile of TAA-APCs post 10-day treatment. MEKi-treated cells expressed VSMCs markers in addition to pericyte markers. N = 5. Images are representative of 1 sample. Scale bar = 50 µm. NG2=Neural-Glial Antigen 2, PDGFRβ= Platelet-Derived Growth Factor Receptor beta, αSMA= Alpha Smooth Muscle Actin.

Compared with the vehicle, PD0325901 significantly suppressed TAA-APC proliferation **(Figure 6A).** Analysis of the secretome revealed that PD0325901-treated TAA-APCs released lower amounts of the anti-angiogenic factor ANGPT-2 **(Figure 6B)**, which was reflected in an improved angiogenic activity of TAA-APCs when co-cultured with AoECs, as observed in an angiogenic network formation assay **(Figure 6C).** Moreover, PD0325901 significantly inhibited TAA-APC motility, as shown in the scratch migration assay **(Figure 6D)**. Interestingly, PD0325901 treatment also inhibited cell sprouting from adventitial explants **(Supplementary Figure 3).** Together, this shows the treatment effectively rescues APC aberrations and shifts the cells towards a reparative and stabilizing phenotype.

**Figure 6.**
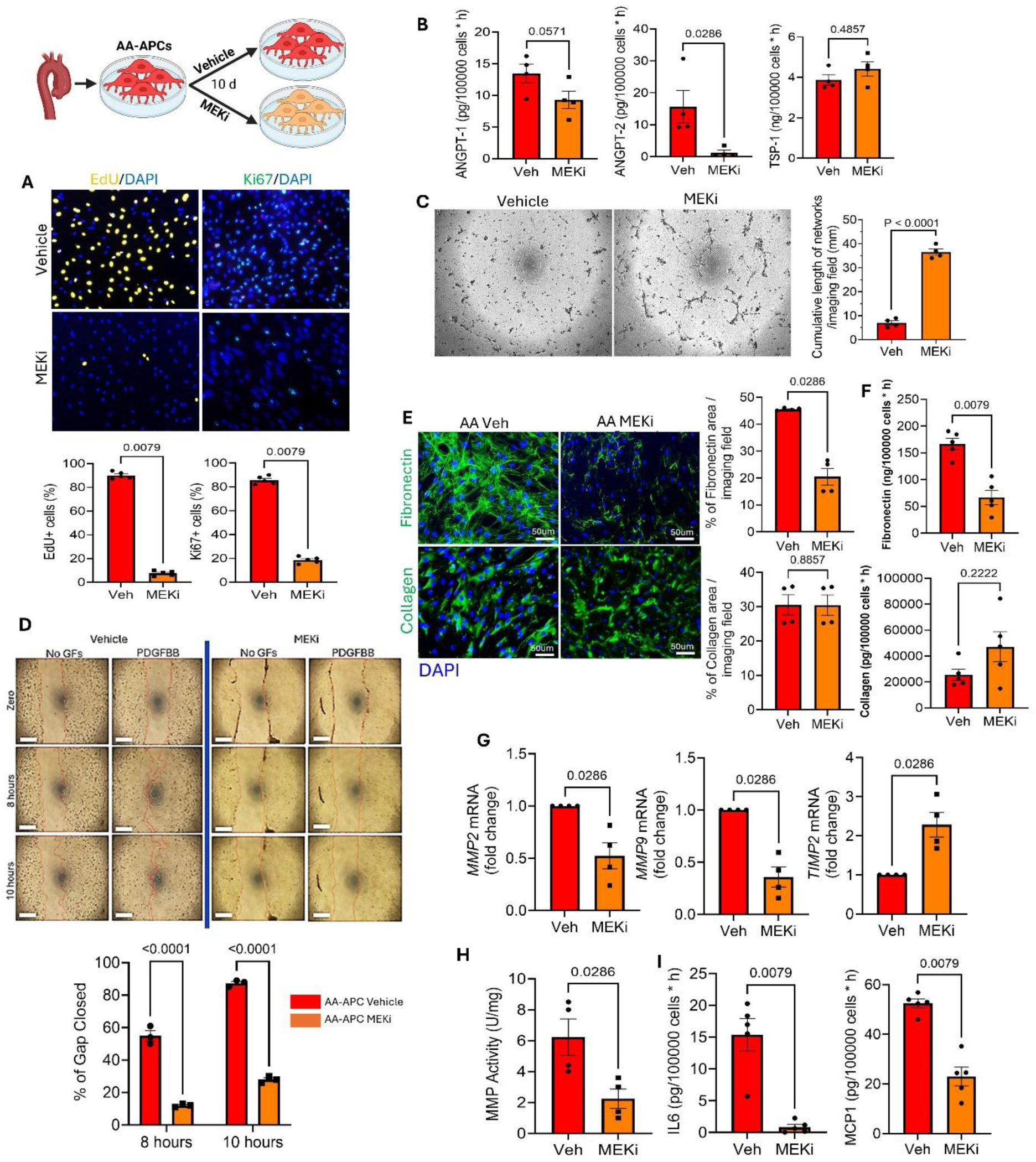
In-vitro PD0325901 treatment restored TAA-APC functional and molecular properties. Comparison of APCs treated for 10 days with either PD0325901 (MEKi) or vehicle. **(A)** Proliferation assays using EdU and Ki67. Images are representative of 1 sample from each group. Yellow/green nuclear fluorescence indicate active proliferating cells. DAPI labels nuclei in blue. Scale bar = 50 µm. N = 5. **(B)** Secreted angiogenic factors in APC conditioned media measured by ELISA. N = 4. **(C)** Angiogenic tube formation assay with MEKi/Vehicle-treated TAA-APCs co-cultured with human aortic ECs (AoECs). Images are representative of 1 sample from each group. N = 4 APCs, n=1 AoECs. **(D)** Wound closure scratch migration assay comparing non stimulated and PDGFBB-stimulated cells. Images were snapped after 8 and 10 hours and compared with time zero. Bar graphs report the percentage of gap closed. N = 3. PDGFBB = platelet-derived growth factor-BB. **(E)** Immunofluorescence images and bar graphs showing fibronectin and collagen (green fluorescence) expression in APCs. DAPI labels nuclei in blue. N = 4. **(F)** Secreted ECM proteins fibronectin and collagen in cell conditioned media. N = 5. **(G)** Expression of *MMP2*, *MMP9* and *TIMP-1* transcripts as assessed by RT-qPCR. N = 4. **(H)** Gelatinase activity fluorescence assay. Results are presented as U/mg. N = 4. **(I)** Secreted proinflammatory cytokines IL-6 and MCP-1 in APC conditioned media. N = 5. All data are presented as individual values and means ± SEM. Analysis by unpaired Mann-Whitney U test.

Furthermore, PD0325901-treated cells produced significantly lower amounts of fibronectin, as assessed using ICC and ELISA **(Figure 6E&F)**. Conversely, treatment did not affect collagen synthesis. PD0325901 conditioning reduced *MMP2* and *MMP9* transcripts in TAA-APCs, along with increased expression of *TIMP-2* **(Figure 6G)**. Importantly, PD0325901-treated TAA-APCs showed reduced gelatinase activity **(Figure 6H).** These effects have positive implications on ECM synthesis and organization.

Finally, the MEKi treatment significantly reduced the levels of pro-inflammatory cytokines MCP-1 and IL-6 in TAA-APCs, indicating a suppression of the inflammatory response **(Figure 6I)**.

All together, these findings suggest that PD0325901 can effectively restore altered properties of AA-APCs by downregulating the MEK/ERK signaling pathway.

### PD0325901 halts aneurysm progression and preserves aortic wall integrity in a mouse model of aortic aneurysm

An *in vivo* study explored the effects of PD0325901 treatment in a C57BL/6J mouse model of aortic aneurysm. After 72 hours from subcutaneous implantation of ANG-II minipumps, mice were given oral MEKi (PD0325901) or vehicle (DMSO) daily for 14 days **(Figure 7A)**. Healthy mice not undergoing any treatment were used as controls. The effective inhibition of ERK1/2 phosphorylation in the aorta of the PD0325901 group was confirmed using western blotting **(Figure 7B).**

**Figure 7.**
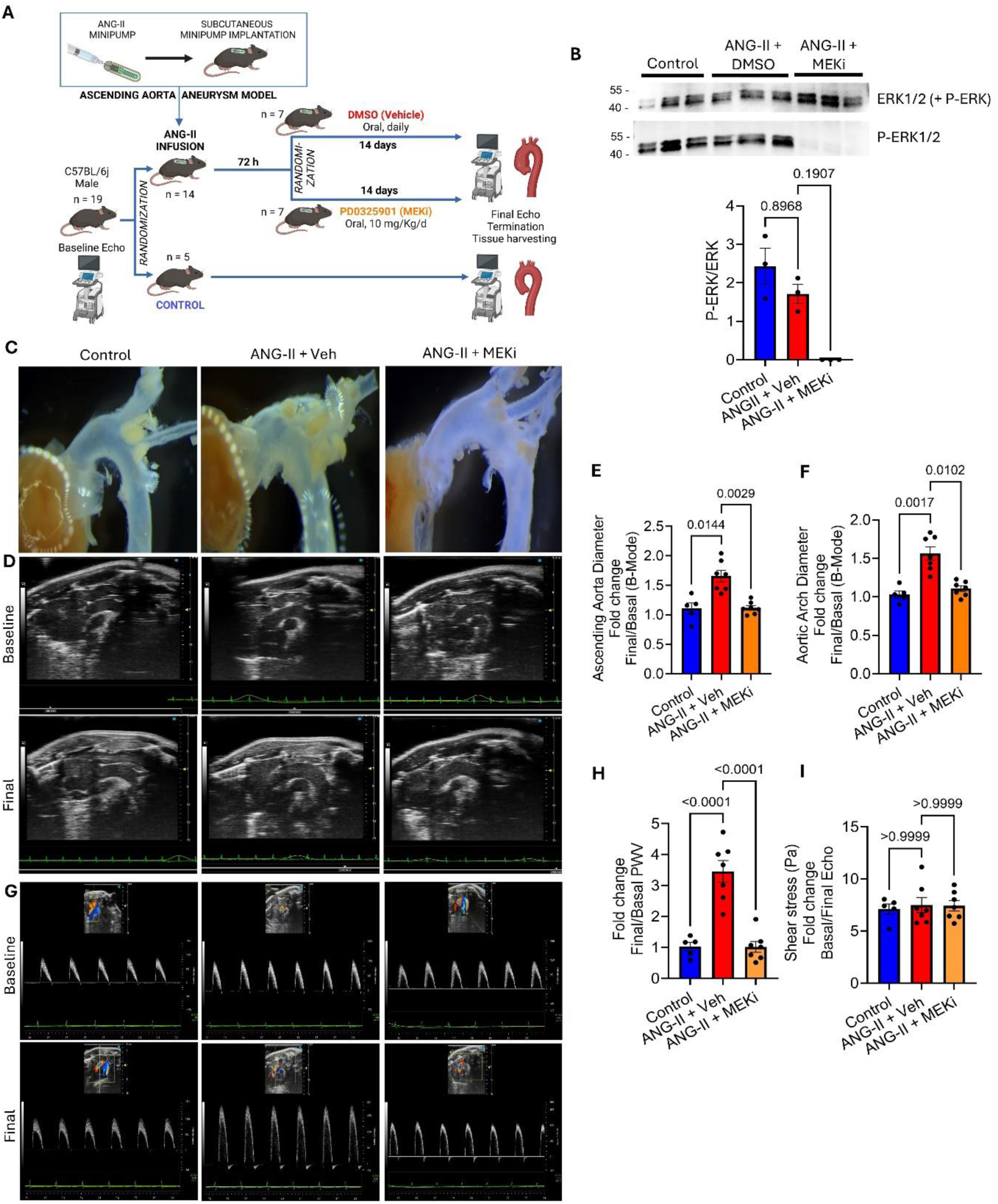
Oral PD0325901 treatment for 14 days halted TAA progression in a murine AA model. **(A)** Cartoon showing the experimental protocol. Mice were started on oral PD0325901 (10 mg/kg/d) or vehicle (DMSO) embedded in jelly 72 hours post ANG-II-minipump implantation and for 14 days. Control mice were included as a reference and did not receive any treatment. **(B)** Western blot showing protein expression for total ERK1/2 and P-ERK in aorta lysates. N = 3 per group. **(C)** Images acquired using a stereo microscope (Leica) showing the mouse aortas after harvest. **(D)** Images of baseline and final echocardiography showing ascending aorta and proximal arch. Images representative of 1 sample from each group. **(E&F)** Inner edge to edge diameters were presented as a fold change of the measurements at the final against basal timepoints. **(G)** PWD images of baseline and final echocardiography representative of 1 sample per group. PWV was calculated using the equation (d-d0)/(T2-T1), where (d-d0)= distance, T1= start time, and T2= end time [PMID: 27824871]. **(H)** PWV calculations were presented as final/basal fold change. **(I)** Oral PD0325901 treatment has no effect on aorta shear stress. Equation used is (wall shear stress (WSS) = 4 * 0.0035 * mean velocity (m/s) / diameter (m)), where (0.0035 Pa.s) is the standard mouse blood viscosity [PMID: 15807389]. In **(E, F, H, I)** N = 5 control, n = 7 vehicle and MEKi. All data are presented as individual values and means ± SEM. All data were analysed by Kruskal-Wallis followed by Dunn’s test.

As previously established ^24,25^, ANG-II treatment induced a dilatation of ascending aorta as assessed by comparison with untreated controls **(Figure 7C)**. Here we focused on the effects of PD0325901 treatment in the TAA group. After 14 days, PD0325901-treated mice showed reduced ascending and arch aorta dilatation compared to vehicle-treated mice **(Figure 7C)**. In the vehicle group, there was an absolute increase in ascending aorta inner diameter from basal to final echo of 0.69±0.06 mm compared with 0.16±0.11 mm in the treated group, P<0.01, **Figure 7D&E**). Consistently, aortic arch diameter increased by 0.57±0.02 mm in final from basal echo in vehicle mice vs. 0.13±0.01 mm in the treatment group, P<0.05, **Figure 7D&F**). Measurement of pulsed wave doppler (PWD) revealed that pulsed wave velocity (PWV), an arterial stiffness index clinically used for risk stratification of TAA, was significantly lower in MEKi-treated mice (with final to basal difference is -2.1 ± 63.23 cm/s) compared to vehicle mice (PWV increased by +497 ± 116 cm/s in final echo compared to basal, P<0.0001) **(Figure 7G&H)**. Shear stress remained unchanged in all groups **(Figure 7I)**. All acquired echocardiography indices are reported in **Supplementary Table 7**. Importantly, no mortality was documented in either TAA groups, suggesting the treatment was safe.

Histological analyses of aortic samples showed that PD0325901 preserved the aortic medial area (P < 0.05, **Figure 8A**) and elastin content (P < 0.05, **Figure 8B**) and inhibited collagen deposition and fibrosis (P < 0.001, **Figure 8C)**, which is in line with reduced stiffness highlighted by the PWV data. Moreover, IHC analysis showed that oral PD0325901 treatment significantly increased the expression of the VSMC marker α-SMA in the media **(Figure 8D)**. In the adventitia, PD0325901 treatment significantly reduced CD45+ inflammatory cell recruitment **(Figure 8E)** as well as adventitial cell apoptosis **(Figure 8F)** with no effect on adventitial PDGFRβ+ cell density **(Figure 8G)**.

**Figure 8.**
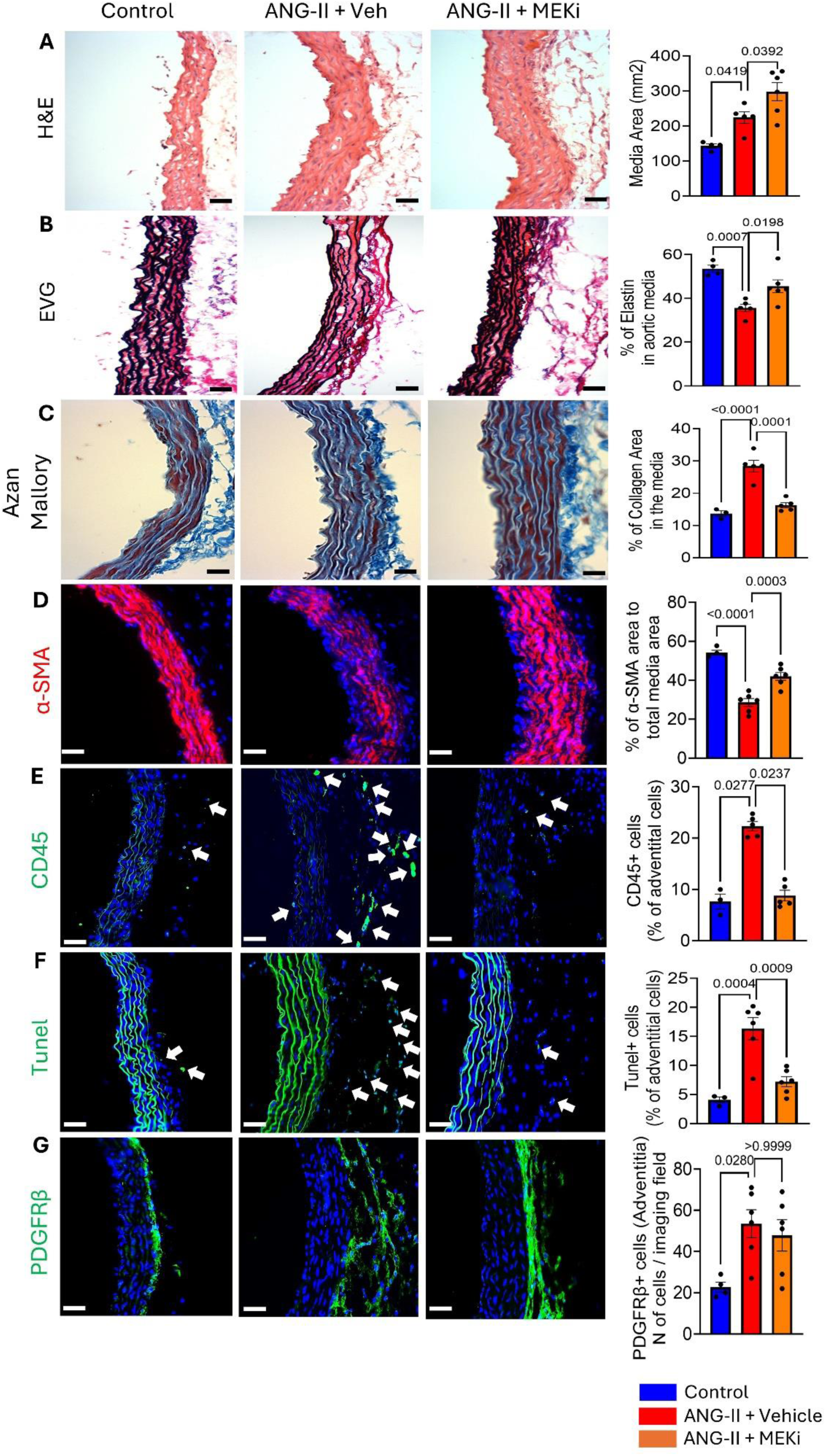
PD0325901 treatment inhibited ANG-II-induced remodelling and enhanced aortic wall integrity. **(A)** Haematoxylin & Eosin images and bar graphs showing aortic medial area. **(B)** EVG staining images, and bar graphs showing elastin medial content. **(C)** Azan Mallory images and bar graphs showing medial fibrosis (collagen content). **(D)** IHC images and bar graphs showing expression of α-SMA (red fluorescence) in medial VSMCs. **(E)** IHC images and bar graphs showing recruitment of CD45+ cells (green). Arrows point to positive cells. **(F)** Fluorescence images and bar graphs showing apoptotic adventitial cells (Tunel+, in green). Arrows point to positive nuclei. **(G)** IHC images and bar graphs showing pericytes (PDGFRβ+ cells) in the adventitia. In all fluorescent images, DAPI was used for nuclei staining (blue). All images are representative of 1 sample from each group. Scale bar = 50 µm. Data are presented as individual values and means ± SEM. Analysis by ordinary one-way ANOVA followed by Dunn’s multiple comparisons test.

According to these data, oral PD0325901 treatment halted the progression of TAA formation, preserved aortic wall integrity, and prevented aortic wall stiffness.

## Discussion

Recent studies have highlighted aortic adventitia as a key site for initial pathological remodeling associated with TAA ^6,7,27^. In this context, our research focused on characterizing aortic APCs and elucidating their role in the development of ascending TAA. We provide robust evidence that APC dysfunction, driven by upregulated MEK/ERK signaling, contributes significantly to ascending TAA progression. Importantly, we demonstrated that this pathological signaling can be modulated using the clinically available, selective MEKi, PD0325901, to rescue APC function and halt TAA progression (mechanism summarized in **Figure 9**).

**Figure 9.**
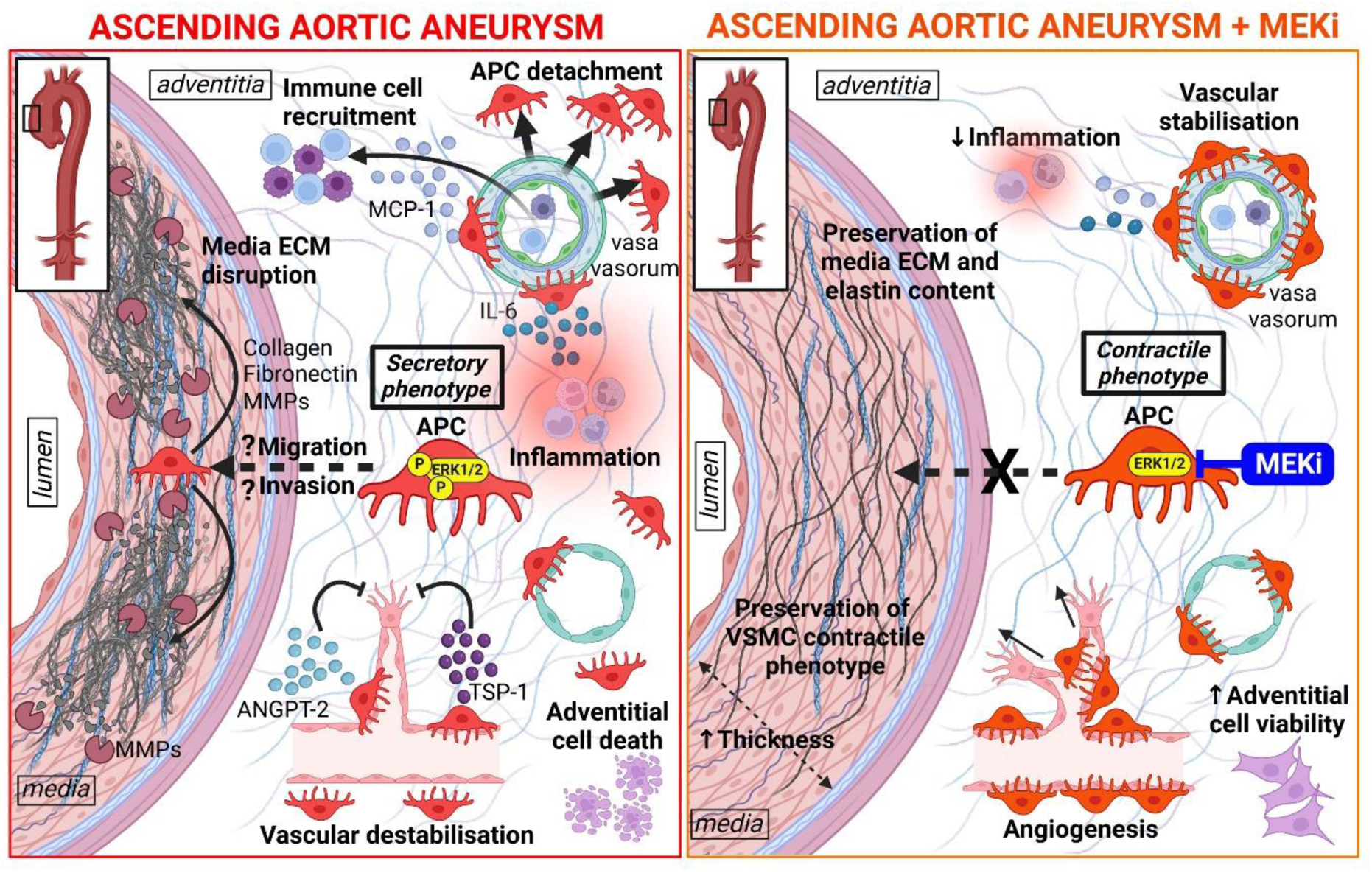
Cartoon summarising the proposed APC-driven mechanism contributing to the pathogenesis of ascending TAA **(left panel)**, and the beneficial effects of MEKi therapy (PD0325901) to rescue APC function and halt TAA progression **(right panel)**. Created in BioRender.com.

Histological analyses of human ascending TAA samples by Billaud et al revealed notable morphometric alterations in the adventitial VV, characterized by reduced density and enlarged vessels with wider lumens and thicker walls ^6^. Our findings corroborated these observations and further identified a significant reduction (by approximately 30%) in VV-associated PDGFRβ+/CD34+ APC coverage compared to control aortas. Furthermore, this reduction was inversely correlated with VV perimeter, lumen diameter and wall thickness (data not shown) suggesting that APC loss contributes directly to VV instability and vascular fragility. Intriguingly, we show that TAA-APCs are more abundant in areas far from the VV, suggesting they detach and migrate away from the VV into the adventitia. Consistently, previous studies showed an increased density of CD34+ progenitor cells in the aortic outer media of TAA patients, suggesting an invasive behavior ^28,29^. Pericytes are essential in regulating endothelial cell stability, vascular tone, and maintaining microvascular integrity ^30^. The observed loss of VV-APC coverage in aneurysm samples mirrors findings in other vascular pathologies, where pericyte loss is associated with increased vascular permeability, hypoxia, and inflammation ^31–33^. In the context of TAA, the associated upregulation of ANGPT-2 and TSP-1 suggests a pathologic phenotype oriented toward destabilizing adventitia vascularization.

Isolated APCs displayed typical pericyte markers (PDGFRβ and NG2) while lacking endothelial and VSMC markers ^13^. Interestingly, there were no antigenic differences between control and aneurysm-isolated APCs, indicating that phenotypic changes in TAA-APCs are driven mainly by altered cellular function rather than antigenic variation. The shift towards a dysfunctional phenotype is induced by aneurysm microenvironmental cues and dysregulated intracellular signaling pathways, including MEK/ERK pathway ^27,34^. The heightened proliferative and migratory capacities of TAA-APCs are linked to vascular destabilization and de-maturation contributing to VV remodeling ^35^. Furthermore, Increased APC migration exacerbates ECM degradation as evidenced by high gelatinase activity and *MMP9* gene expression. This, in addition to the dysregulated ECM protein formation, weakens the structural integrity of the aortic wall, rendering it more susceptible to hemodynamic stress and further dilatation. Moreover, the increased levels of pro-inflammatory cytokines, including MCP-1 and IL-6, create a perpetuating inflammatory loop, recruiting immune cells to the adventitia and further destabilizing the vascular microenvironment ^25,36^.

The impaired angiogenic support provided by TAA-APCs contributes to the pathology. As key regulators of vascular stability, pericytes typically facilitate angiogenesis and maintain endothelial cell integrity ^37,38^. However, the inability of TAA-APCs to form stable networks with AoECs, associated with disrupted angiogenic factors secretion, indicates a loss of function critical for adventitia VV homeostasis. This disruption contributes to the reduced VV density observed in aneurysm tissue, which, in turn, impairs oxygen and nutrient delivery to the outer layers of the aorta ^6^. Hypoxia within the adventitia can further stimulate pathological angiogenesis, perpetuating VV remodeling and promoting aneurysm expansion ^6,7^. Importantly, the expanding aortic wall should be compensated by increased angiogenic activity in the early TAA stage. However, APC dysfunction and angiogenic impairment may result in the low angiogenic profile found in the late TAA stage (i.e., surgical samples collected).

*In vitro* studies showed that the selective MEKi, PD0325901, restored the contractile phenotype of TAA-APCs and demonstrated therapeutic effects on their functional properties. This confirms that MEK/ERK pathway hyperactivation, the hallmark signaling in AA ^18,19^, is a central driver of APC dysfunction in TAA, and its inhibition can effectively reprogram APCs toward a physiologic phenotype. The induction of VSMC markers, including α-SMA, Calponin, and SM-MHC, highlights the APC shift from synthetic to contractile phenotype. This phenotypic switch supports VV integrity and limits adventitial remodeling by obligating cellular differentiation into a reparative phenotype.

PD0325901 treatment also normalized TAA-APCs secretome, reducing the levels of ANGPT-2, MCP-1, and IL-6, and the balance of matricellular proteins. These factors are key in disrupting angiogenesis, driving inflammation, and promoting ECM degradation. Translating the *in vitro* data into a clinical scenario, we propose that MEKi would restore adventitia homeostasis by multiple protective mechanisms. These properties could help prevent TAA enlargement to the ultimate stage when only surgery can work.

To challenge this hypothesis, the therapeutic efficacy of PD0325901 was further validated in a mouse TAA model ^24,25^ induced by the vasoconstrictor Ang-II, which is a potent stimulus for MEK/ERK, c-Jun N-terminal protein kinase (JNK), and p38 ^39,40^. Our results provide strong evidence for the potential of PD0325901 in preventing the progression of TAA and improve aortic wall integrity. By effectively inhibiting P-ERK, the treatment addresses a key signaling driver of TAA progression, significantly improving the aortic wall’s structural and functional parameters. Echocardiographic findings showing reduced diameters of the ascending aorta and aortic arch highlight the ability of PD0325901 to attenuate aneurysmal dilation. Moreover, the accompanying reduction in PWV, a strong indicator of wall stiffness ^41,42^, reflects improved aortic compliance, and confirms that the treatment not only halts aneurysm expansion but also restores the biomechanical properties of the aortic wall, which are typically compromised in TAA. Moreover, the non-significant changes in wall shear stress refer to the targeted effect of PD0325901 on preserving aortic wall integrity with no effect on hemodynamics.

Histological evidence further supports these findings, demonstrating a multifaceted therapeutic impact of PD0325901 treatment on the aortic wall. Increased α-SMA expression in the media confirms enhanced VSMC stability, while improved elastin content and reduced fibrosis indicate the restoration of ECM integrity. The preservation of medial area underscores the potential of PD0325901 to counteract the pathological medial remodeling and aortic wall thinning, which are hallmarks of aneurysm development and progression ^43,44^. Furthermore, the observed reduction in adventitial inflammation and apoptosis has significant therapeutic importance, as chronic inflammation and cell death are major contributors to TAA development ^25,45,46^. Finally, the non-significant difference in terms of adventitia PDGFRβ+ cell density between vehicle and MEKi groups could be due to the starting time of oral MEKi treatment (72h post-ANG-II infusion), when the APC density achieved at this stage might not be later affected by treatment. By preserving PDGFRβ+ APC density and – importantly – restoring their physiological function, PD0325901 supports adventitial integrity, further reinforcing the aorta’s overall structural resilience. Notably, the treatment achieves these effects without compromising the viability of APCs or manifesting safety concerns for mice. However, further studies are required to delineate the effects of PD0325901 on TAA over a longer time frame.

### Conclusion

Our findings suggest that PD0325901 effectively modulates pathological MEK/ERK signaling in APCs, restoring their contractile phenotype and normalizing their function. This therapeutic approach has the potential to halt aneurysm growth and restore critical structural and functional properties of the aortic wall, making it a promising candidate for future therapeutic development.

### Study limitations

1. The characterization of VV and pericytes in human TAA was performed on surgical samples at the late stage of the disease. We could not perform similar analyses on early TAA samples (when surgery is not indicated), which would have helped us dissect the cellular and molecular evolution of the disease. The in vivo study, however, although performed in a different species and with TAA induction by high doses of ANG-II, confirms that pericytes and ERK signaling could be therapeutic targets to prevent TAA expansion.
2. Given the distance in species, the translational value of our data should be considered with caution. Moreover, additional pharmacokinetic and safety studies are needed to determine the optimal treatment schedule and safety.

## Acknowledgements

Drawings were created using Biorender.com.

## Data Availability

The data that support the findings of this study are available from the corresponding authors upon request.

## Authors contribution

KAKM, EA, and VVA conducted the experiments and acquired the data.

KAKM and EA analyzed the data and drafted the manuscript.

KAKM, EA and PM interpreted the data.

EA, GDA, and PM supervised the research.

PM contributed to the research conception and design, and critically read the final manuscript.

GDA, EMA, and CR recruited patients and provided human samples.

AE and AG secured funding.

All authors approved the authorship order and the final version of the manuscript for publication.

## Supplemental Material

Tables S1–S7 Figure S1, S2

