## Supplementary figures and tables for "Molecular reprogramming of adventitial pericytes by a selective MEK inhibitor halts the progression of thoracic aortic aneurysm"

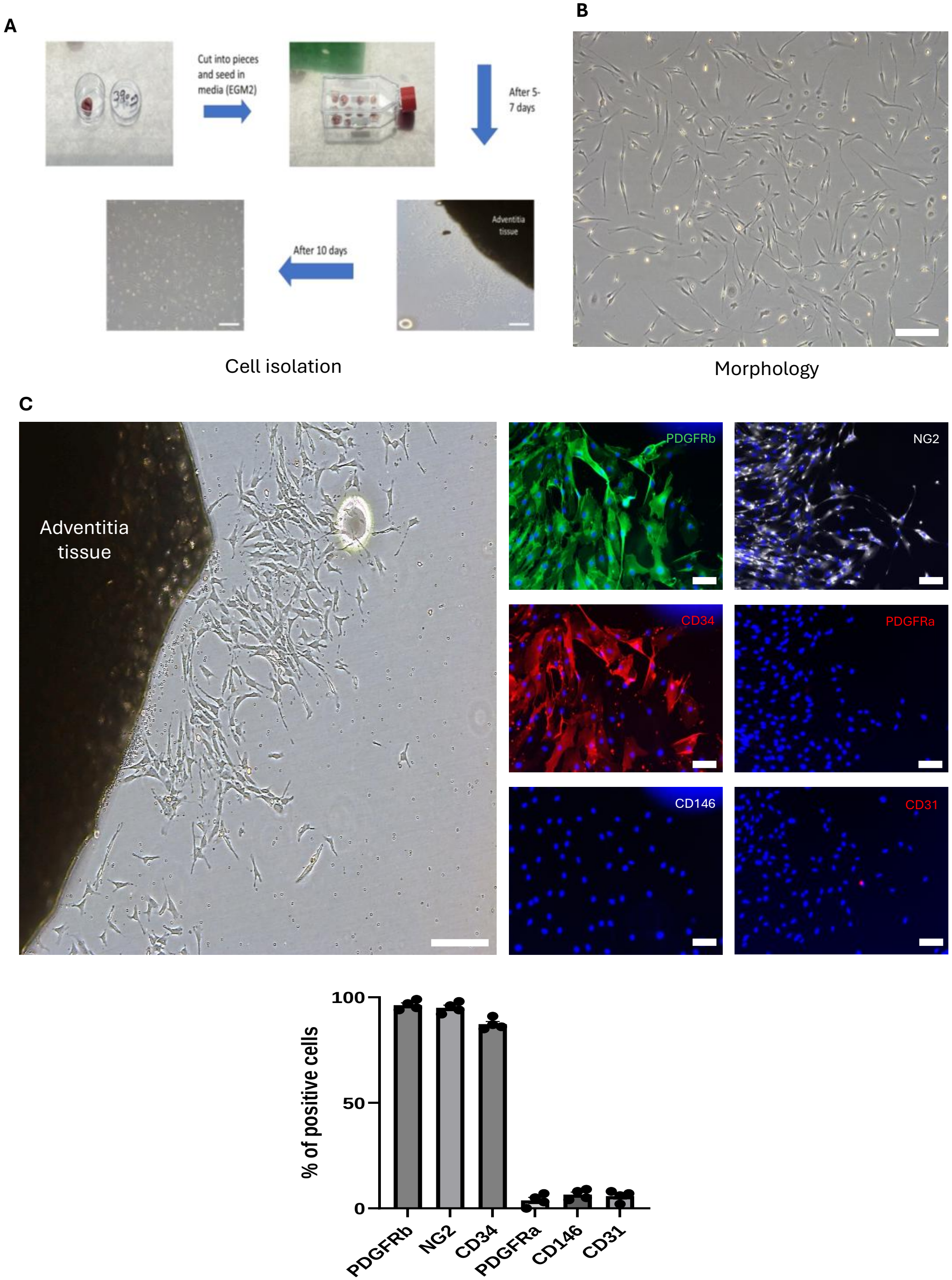


**Suppl. Figure 1**

**Suppl. Figure 1. Isolation and antigenic characterization of APCs. (A)** Explant outgrowth method. Adventitia samples were cut into small pieces, cultured in tissue culture flasks with full EGM2 media and incubated in an incubator (37 C, 5% $CO_2$). Cells started to sprout after 5-7 days, pieces were removed, and cells expanded. **(B)** Brightfield optical image of sprouting cells at passage 1, showing morphology of pericytes (i.e., spindle shape with small central nuclei). **(C)** Immunofluorescence images showing the antigenic profile of the sprouting cells at passage zero. N = 4. Images are representative of 1 sample. Data are presented as individual values and means ± SEM.

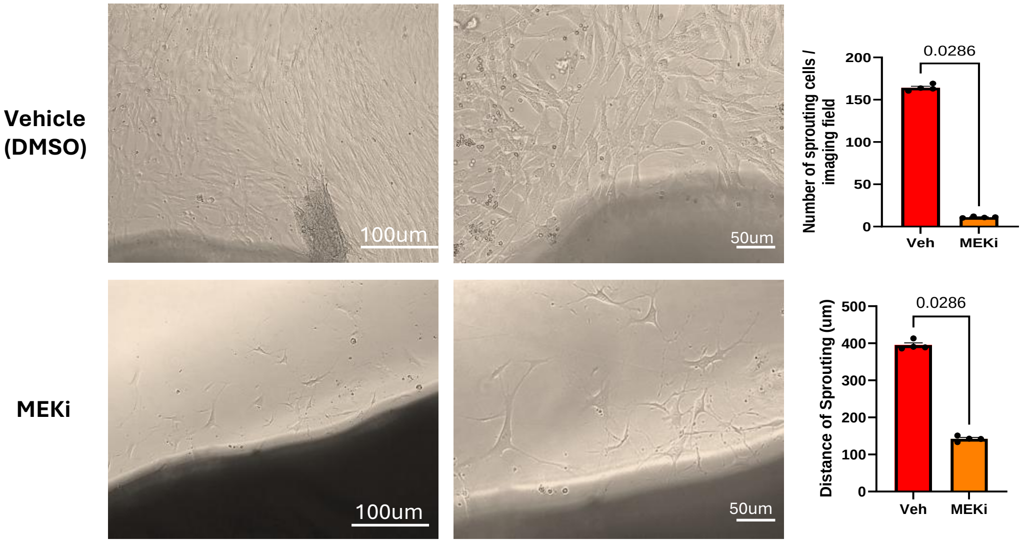


**Suppl. Figure 2. Ex-vivo study to assess the effect of PD0325901 on APC sprouting from adventitia explants.** Brightfield optical images showing cell sprouting from explants incubated with the MEKi (250 nM) or Vehicle (DMSO) for 10 days. MEKi significantly inhibited APC sprouting and migration from tissue. N = 4. Images are representative of 1 sample. Data are presented as individual values and means+/-SEM. Analysis by unpaired Mann-Whitney U test.

**Suppl. Figure 2**

### Supplementary Tables

| Category | Cell line | Surgery | Age (years) | Gender | Asc. Aorta diameter | Experiments |
|---|---|---|---|---|---|---|
| Control | APC 1 | CABG | 57 | Male | 1.8 cm | Histology, IHC, ICC, Doubling time, Migration, ELISA, Angiogenesis assay, Gelatinase activity, RT-qPCR, WB |
| | APC 2 | CABG | 63 | Male | 2.0 cm | Histology, IHC, ICC, RT-qPCR, ELISA, Doubling time, Migration |
| | APC 3 | CABG | 62 | Male | 1.9 cm | Histology, IHC, ICC, Angiogenesis assay, RT-qPCR, WB, ELISA, Gelatinase activity, WB |
| | APC 4 | CABG | 62 | Male | 1.9 cm | ICC, RT-qPCR, Angiogenesis assay, Doubling time, Migration, ELISA |
| | APC 5 | CABG | 68 | Female | 2.1 cm | ICC, RT-qPCR, WB, Gelatinase activity, Doubling time, ELISA |
| | APC 6 | CABG | 57 | Female | 2.7 cm | Histology, IHC, RT-qPCR, Doubling time, ELISA, Migration, WB, Angiogenesis assay, Gelatinase activity |

**Table 1. Control Aortas.** Patients and samples characteristics, and list of experiments in which the extracted cells were used.

| Category | Cell line | Surgery | Age (years) | Gender | Asc. Aorta diameter | Experiments |
|---|---|---|---|---|---|---|
| Aortic Aneurysm | AAPC 1 | Asc. AA graft replacement | 79 | Female | 5.5 cm | **Histology, IHC, ICC, Migration, ELISA, Angiogenesis assay, Gelatinase activity, RT-qPCR, WB** |
| | AAPC 2 | Asc. + Arch AA graft replacement | 54 | Female | 5.3 cm | **Histology, IHC, ICC, RT-qPCR, ELISA, Doubling time, Migration, in-vitro MEKi** |
| | AAPC 3 | Root + Asc. AA graft replacement | 61 | Male | 5.8 cm | **Histology, IHC, ICC, Angiogenesis assay, RT-qPCR, WB, ELISA, Gelatinase activity, WB, in-vitro MEKi** |
| | AAPC 4 | Asc. AA graft replacement | 73 | Male | 5.6 cm | **Histology, IHC, ICC, RT-qPCR, Angiogenesis assay, Doubling time, Migration, ELISA, in-vitro MEKi** |
| | AAPC 5 | Root + Asc. AA graft replacement | 56 | Female | 6.1 cm | **ICC, RT-qPCR, WB, Gelatinase activity, Doubling time, ELISA** |
| | AAPC 6 | Root + Asc. + Arch AA graft replacement | 62 | Male | 5.8 cm | **Histology, IHC, RT-qPCR, Doubling time, ELISA, WB, Angiogenesis assay, Gelatinase activity, in-vitro MEKi** |
| | AAPC 7 | Asc. AA graft replacement | 68 | Male | 5.7 cm | **ICC, ELISA, Angiogenesis assay, Gelatinase activity, RT-qPCR, in-vitro MEKi** |
| | AAPC 8 | Root + Asc. AA graft replacement | 71 | Female | 6.2 cm | **ICC, Migration, ELISA, Angiogenesis assay, RT-qPCR.** |
| | AAPC 9 | Asc. + Arch AA graft replacement | 64 | Male | 6.5 cm | **Histology, IHC, ICC, Migration, Doubling time, Angiogenesis assay, RT-qPCR.** |

**Table 2. Aneurysm Aortas.** Patients and samples characteristics, and list of experiments in which the extracted cells were used.

| Target | Host | Dilution | Company & CAT# |
|---|---|---|---|
| PDGFRβ (anti-human) | Goat | 1:50 | R&D AF385 |
| PDGFRβ (anti-mouse) | Goat | 1:50 | R&D AF1042 |
| αSMA | Mouse (conjugated) | 1:400 | Dako M0851 |
| CD34 | Mouse | 1:100 | Dako M7165 |
| CD31 | Rabbit | 1:20 | Abcam Ab28364 |
| CD45 | Rat | 1:500 | NovusBio NB100-77417 |

**Table 3**. Primary antibodies used for immunohistochemistry on human and mouse tissues.

| Primary Ab | Host | Company & CAT# | Dilution | Triton-X100 (permeabilization) |
|---|---|---|---|---|
| CD34 | Mouse | Dako M7165 | 1:100 | No |
| CD31 | Rabbit | Abcam Ab28364 | 1:50 | No |
| CD146 | Rabbit | Abcam Ab75769 | 1:100 | Yes |
| NG2 | Rabbit | Millipore AB5320 | 1:50 | No |
| PDGFRβ | Goat | R&D AF385 | 1:50 | No |
| PDGFRα | Mouse | Santa Cruze sc-398206 | 1:100 | Yes |
| SM α-Calponin | Rabbit | Abcam Ab46794 | 1:100 | Yes |
| α-SM-MHC11 | Rabbit | Abcam Ab53219 | 1:50 | Yes |
| Smoothelin B | Rabbit | Santa Cruz sc-28562 | 1:100 | Yes |
| α-SMA | Mouse | Dako M0851 | 1:100 | Yes |
| Transgelin (SM22α) | Mouse | Santa Cruz sc-271719 | 1:50 | Yes |
| Vimentin | Rabbit | Abcam Ab92547 | 1:500 | Yes |
| Ki67 | Rabbit | Abcam, ab16667 | 1:500 | Yes |
| P-ERK (Thr202/ Tyr204) | Rabbit | Cell Signalling, #4370 | 1:200 | Yes (Methanol) |

**Table 4**. Primary antibodies used for immunocytochemistry in human cells.

| Tested Gene | ID |
|---|---|
| ***MMP 2*** | HS01548727 |
| ***MMP 9*** | HS00234579 |
| ***TIMP 2*** | HS00234278 |
| ***GAPDH* (Housekeeping)** | HS02786624 |

**Table 5**. List of RT-qPCR probes.

| Target | Host | Dilution | Company & CAT# |
|---|---|---|---|
| **ERK 1/2 (p44/42 MAP)** | Rabbit | 1:1000 | Cell Signalling, #4695 |
| **P-ERK (Thr202/ Tyr204)** | Rabbit | 1:2000 | Cell Signalling, #4370 |

**Table 6**. List of antibodies used for Western Blot.

| | Control | | Vehicle | | MEKi | |
|---|---|---|---|---|---|---|
| Echo indices | Basal | Final | Basal | Final | Basal | Final |
| Ascending diameter | **1.25 ± 0.2** | **1.37 ± 0.1** | **1.1 ± 0.17** | **1.8 ± 0.11** | **1.3 ± 0.06** | **1.5 ± 0.18**** |
| Ascending area | **3.1 ± 0.7** | **3.5 ± 0.5** | **2.8 ± 0.65** | **6.4 ± 0.65** | **2.75 ± 0.49** | **5.1 ± 0.85** |
| Arch diameter | **1.21 ± 0.13** | **1.23 ± 0.03** | **1.07 ± 0.17** | **1.64 ± 0.16** | **1.27 ± 0.09** | **1.4 ± 0.10*** |
| Arch area | **3.3 ± 0.5** | **3.5 ± 0.3** | **2.76 ± 0.83** | **5.02 ± 0.66** | **3.04 ± 0.32** | **4.73 ± 0.54** |
| PWV | **202.6 ± 45.6** | **197.3 ± 40.8** | **209.99 ± 32.61** | **706.99 ± 148.12** | **246.84 ± 47.58** | **244.71 ± 110.81******** |
| Shear stress | **6.9 ± 1.2** | **7.1 ± 1.1** | **9.05 ± 2.16** | **7.49 ± 1.93** | **6.1 ± 1.48** | **7.41 ± 1.35** |

**Table 7. Echocardiography indices in the three groups of mice: control vehicle, and MEKi.** Echocardiography aortic measurements were performed before ANG-II minipump implantation (basal) and after 14 days of oral treatment with vehicle or PD0325901 (MEKi), control mice received oral jelly. Control (n=5), Vehicle (n=7), and MEKi (n=7). Values are means and SD, * P<0.05 , ** P<0.01 , and **** P<0.0001 vs. Final vehicle.
